# Turnover number predictions for kinetically uncharacterized enzymes using machine and deep learning

**DOI:** 10.1101/2022.11.10.516024

**Authors:** Alexander Kroll, Xiao-Pan Hu, Nina A. Liebrand, Martin J. Lercher

## Abstract

The turnover number *k*_cat_, a measure of enzyme efficiency, is central to understanding cellular physiology and resource allocation. As experimental *k*_cat_ estimates are unavailable for the vast majority of enzymatic reactions, the development of accurate computational prediction methods is highly desirable. However, existing machine learning models are limited to a single, well-studied organism, or they provide inaccurate predictions except for enzymes that are highly similar to proteins in the training set. Here, we present TurNuP, a general and organism-independent model that successfully predicts turnover numbers for natural reactions of wild-type enzymes. We constructed model inputs by representing complete chemical reactions through difference fingerprints and by representing enzymes through a modified and re-trained Transformer Network model for protein sequences. TurNuP outperforms previous models and generalizes well even to enzymes that are not similar to proteins in the training set. Parameterizing metabolic models with TurNuP-predicted *k*cat values leads to improved proteome allocation predictions. To provide a powerful and convenient tool for the study of molecular biochemistry and physiology, we implemented a TurNuP web server at https://turnup.cs.hhu.de.

## Introduction

The turnover number *k*_cat_ is the maximal rate at which one activate site of an enzyme converts molecular substrates into products. *k*_cat_ is a central parameter for quantitative studies of enzymatic activities, and is of key importance for understanding cellular metabolism, physiology, and resource allocation. In particular, comprehensive sets of *k*_cat_ values are essential for metabolic models that consider the cost of producing or maintaining enzymes^1–9^, a prerequisite for accurate simulations of cellular physiology and growth^10^. Currently, no high-throughput experimental assays exist for *k*_cat_, and experiments are both time consuming and expensive. Thus, *k*_cat_ estimates are unavailable for most reactions; even for *Escherichia coli*, arguably the biochemically best-characterized organisms, *in vitro k*_cat_ is known for only ∼ 10% of all enzyme-catalyzed reactions^11^. In genome-scale kinetic models of cellular metabolism, this issue is typically addressed by either sampling missing *k*_cat_ values or fitting them to large datasets^7,8,12,13^. However, these techniques typically result in inaccurate results, and fitted *k*_cat_ values bear little relationship to known *in vitro* estimates^7,12,13^.

Recent advances in artificial intelligence have put the computational prediction of unknown *k*_cat_ values from *in vitro* training data into reach, and two recent publications have explored this possibility. Heckmann et al.^14^ developed a *k*_cat_ prediction model for enzymes in *E. coli*. The model relies on detailed, expertcrafted input features such as enzyme active site properties, metabolite concentrations, experimental conditions, and reaction fluxes calculated through flux balance analysis (FBA)^15^. It achieved a coefficient of determination *R*^2^ ≈ 0.34 on an independent test set. However, the complete, detailed input information is only available for a small subset of enzymatic reactions even in *E. coli*, limiting the applicability of this approach. A deep learning model that requires less detailed input features, DLKcat, was recently developed by Li et al.^16^. DLKcat predicts *k*_cat_ using information about the enzyme’s amino acid sequence and about one of the reaction’s substrates, ignoring other reaction details such as products and co-substrates. In practical applications, *k*_cat_ predictions are most important when no experimental measurements for closely related enzymes are available, and hence general prediction models should generalize well to such cases. However, while DLKcat can in principle be applied to any enzymatic reaction, its predictions become misleading for enzymes not similar to those in the training set, as we demonstrate below.

Here, we present a general machine and deep learning approach for predicting *in vitro k*_cat_ values for natural reactions of wild-type enzymes. In contrast to previous approaches, we represent chemical reactions through numerical fingerprints that consider the complete set of substrates and products of a reaction. To capture the enzyme properties, we use fine-tuned state-of-the-art protein representations as additional model inputs (**Figure 1**). We created these enzyme representations using Transformer Networks, deep neural networks for sequence processing, which were trained with millions of protein sequences^17^. It has been shown for various prediction tasks that Transformer Networks outperform protein representations created with convolutional neural networks (CNNs)^18,19^, which were used in previous models for predicting enzyme turnover numbers^16^.

**Figure 1.**
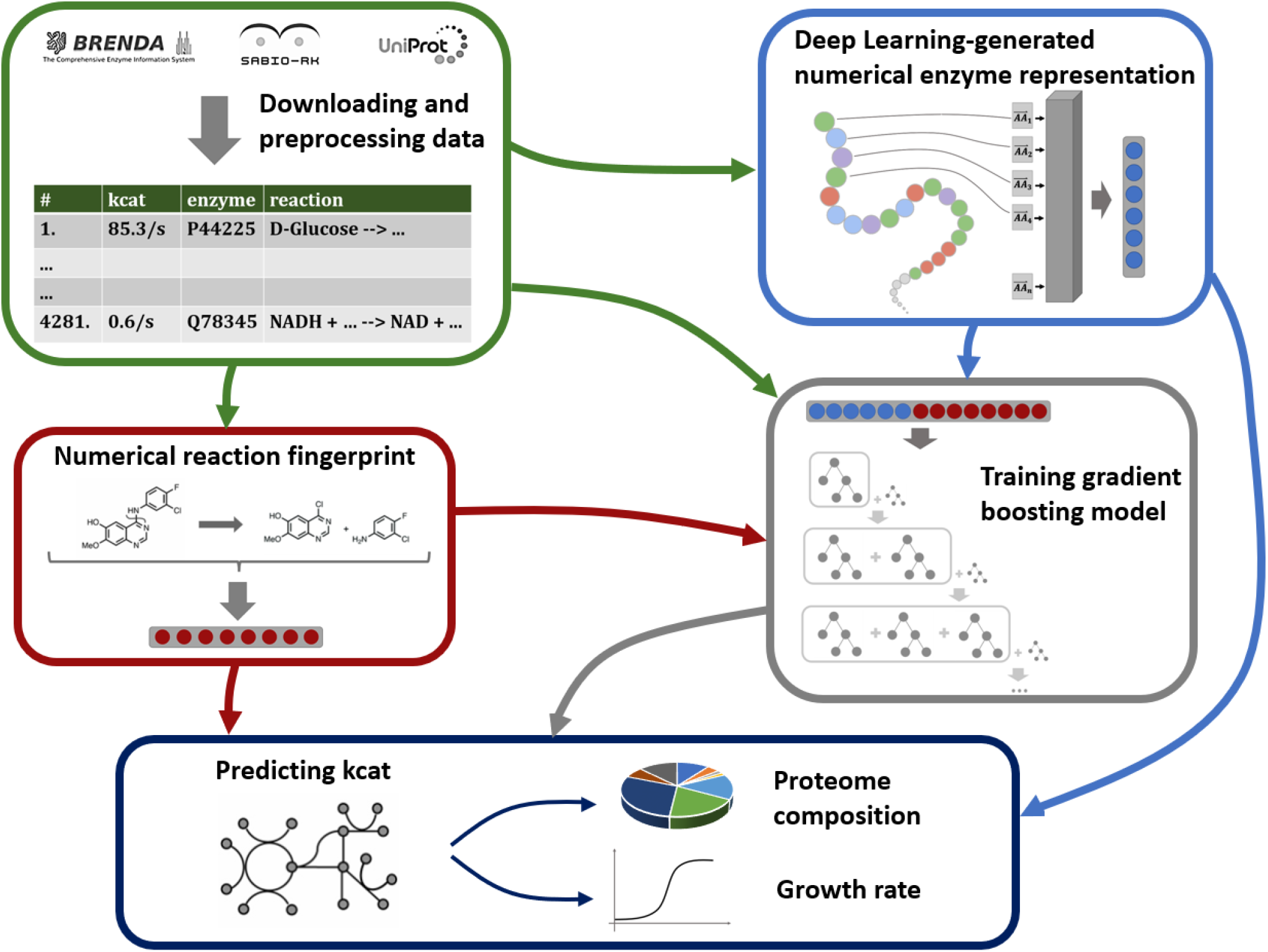
Machine learning model to predict *k*_cat_ from numerical enzyme representations and reaction fingerprints. Experimentally measured *k*_cat_ values are downloaded from three different databases. Enzyme information is represented with numerical vectors obtained from natural language processing (NLP) models that use the linear amino acid sequence as their input. Chemical reactions are represented using integer vectors. Concatenated enzyme-reaction representations are used to train a gradient boosting model to predict *k*_cat_. After training, the fitted model can be used to parameterize metabolic networks with *k*_cat_ values.

Our resulting Turnover Number Prediction model – TurNuP – outperforms both previous methods for predicting *k*_cat_ ^14,16^. We show that TurNuP generalizes well even to enzymes with *<* 40% sequence identity to proteins in the training set. Using genome-scale, enzyme-constrained metabolic models for different yeast species^16^, we demonstrate that parameterizations with TurNuP *k*_cat_ predictions lead to improved proteome allocation predictions. To facilitate widespread use of the TurNuP model, we not only provide a Python function for large-scale *k*_cat_ calculations by bioinformaticians, but we also built an easy-to-use web server that requires no specialized software (turnup.cs.hhu.de).

## Results

### Obtaining training and test data

We compiled a dataset that connects *k*_cat_ measurements with the corresponding enzyme sequences, reactant IDs, and reaction equations. The underlying data is derived from the three databases BRENDA^20^, UniProt^21^, and Sabio-RK^22^. Our aim was to build a turnover number prediction model for natural reactions of wild-type enzymes. We hypothesized that we do not have enough data to train a model to predict the catalytic effect of enzyme mutations or to predict the *k*_cat_ value of non-natural enzyme-reaction pairs, which have not been shaped by natural selection. Hence, we removed all data points with non-wild-type enzymes and all non-natural reactions (see Methods, “Data preprocessing”). We removed redundancy by deleting data that was identical to other data points in the set, and we excluded points with incomplete reaction or enzyme information. We also removed 55 outliers with unrealistically low or high measurements, i.e., reported *k*_cat_ values that are either very close to zero (*<* 10^*−*2.5^*/s*) or that are unreasonably high (*>* 10^5^*/s*)^23^. If multiple different *k*_cat_ values existed for the same enzyme-reaction pair, we took the geometric mean across these values.

This resulted in a final dataset with 4 271 data points, comprising 2 977 unique reactions and 2 827 unique enzymes (for more details on data preprocessing, see Methods). We log_10_-transformed all *k*_cat_ values to obtain a target variable with an approximately Gaussian distribution (**Figure S1**). We split the dataset into 80% training data and 20% test data in such a way that enzymes with the same amino acid sequence would not occur both in the training and in the test set. We further split the training set into 5 disjoint subsets to perform 5-fold cross validations (CVs) for hyperparameter optimization of our machine learning models. To challenge our models to learn to predict *k*_cat_ of enzymes without kinetically characterized close homologs, the cross validation sets were constructed such that no two subsets contained enzymes with identical amino acid sequences.

### Numerical reaction fingerprints alone lead to reasonable *k*_cat_ predictions

The *k*_cat_ value of an enzyme-catalyzed reaction depends strongly on the catalyzing enzyme, but also on the chemical reaction itself. To integrate reaction information into our machine learning model, we used numerical reaction fingerprints. We compared the performance of two different types of such representations, structural and difference fingerprints (calculated with the Python package Chem from RDKit^24^).

To create structural reaction fingerprints, one first calculates for each substrate and each product a 1 638-dimensional binary molecular fingerprint, designed to encode structural information of small molecules. The bit-wise OR-function is then applied to all substrate fingerprints and separately to all product fingerprints, resulting in two 1 638-dimensional binary vectors with molecular information about the substrates and about the products, respectively. These two vectors are concatenated, providing a 3 276-dimensional binary vector with structural information about the reaction^24^ (**Figure 2a**).

**Figure 2.**
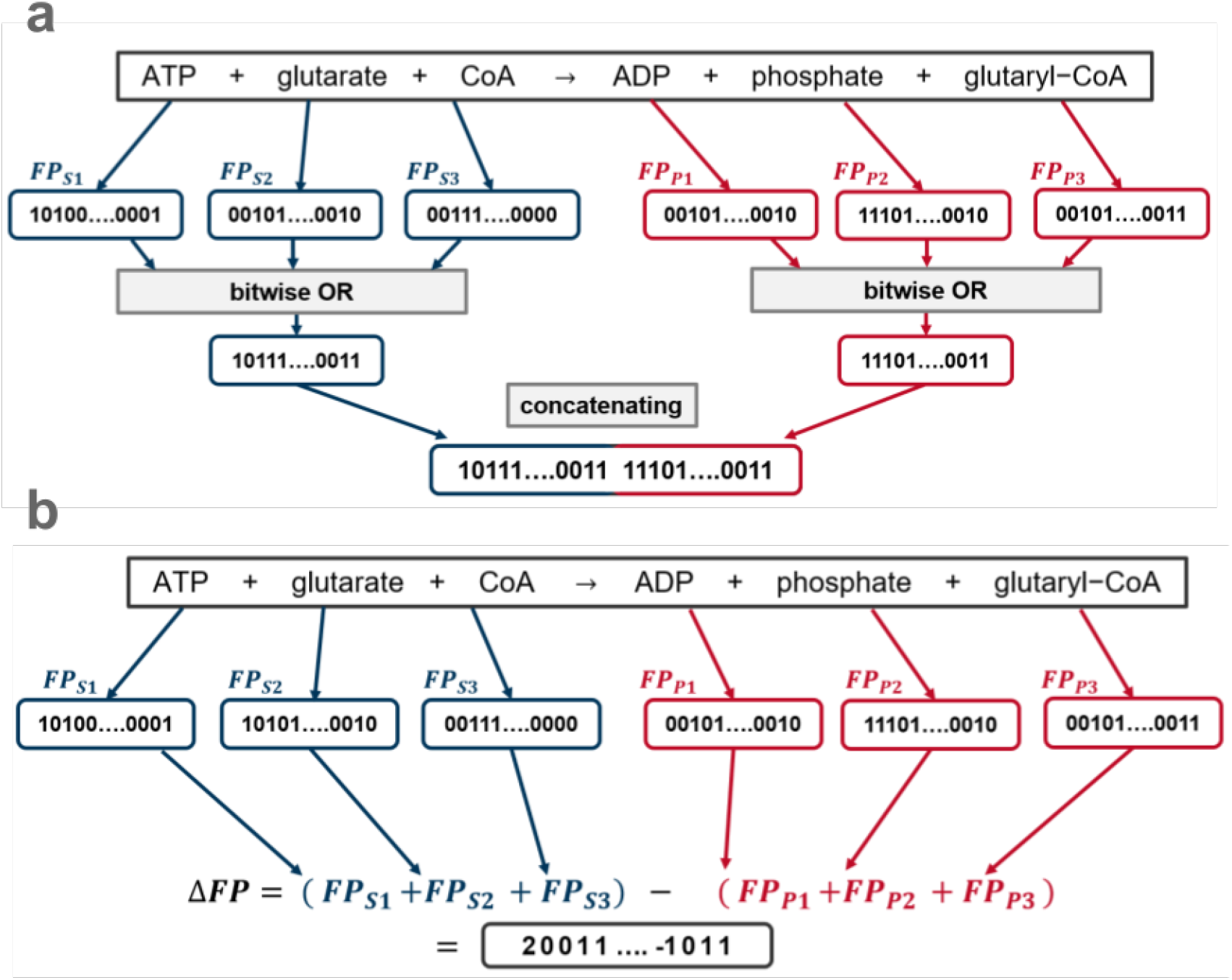
Calculation of reaction fingerprints for an exemplary reaction. **(a)** Structural reaction fingerprints. Binary molecular fingerprints are calculated for each substrate and each product. The bitwise OR-function is applied to all substrates and also to all products. The resulting substrate and the resulting product vector are then concatenated. **(b)** Difference reaction fingerprints. Binary molecular fingerprints are calculated for each substrate and each product. All substrate fingerprint vectors are summed, and the same is done for all product fingerprint vectors. To create the difference fingerprint, the resulting product vector is subtracted from the substrate vector.

The calculation of difference reaction fingerprints starts with a different, 2 048-dimensional binary fingerprint for each substrate and each product. All substrate fingerprint vectors are summed to provide a single substrate vector, and all product fingerprint vectors are summed to provide a single product vector. This product fingerprint is then subtracted from the substrate fingerprint, resulting in a 2 048-dimensional reaction fingerprint with positive and negative integers^25^ (**Figure 2b**).

To test how well the reaction fingerprints alone can predict the turnover numbers of enzyme-catalyzed reactions, we trained two gradient boosting models to predict *k*_cat_, each with one of the reaction fingerprints as the only input. We performed a 5-fold CV with a random grid search for hyperparameter optimization for both models. After hyperparameter optimization, we chose the set of hyperparameters with the highest coefficient of determination *R*^2^ across CV sets, and we re-trained each model with its best hyperparameters on the whole training set. On the test set, the resulting model with structural reaction fingerprints as its inputs achieves a coefficient of determination *R*^2^ = 0.31, a mean squared error *MSE* = 0.99, and a Pearson correlation coefficient *r* = 0.56 on the test set. The model with difference reaction fingerprints achieves slightly improved results, with *R*^2^ = 0.34, *MSE* = 0.95, and *r* = 0.60 on the test set (**Figure 3**). Thus, a model based on chemical reaction information alone can already predict about a third of the variation in *k*_cat_ across enzyme-catalyzed reactions.

**Figure 3.**
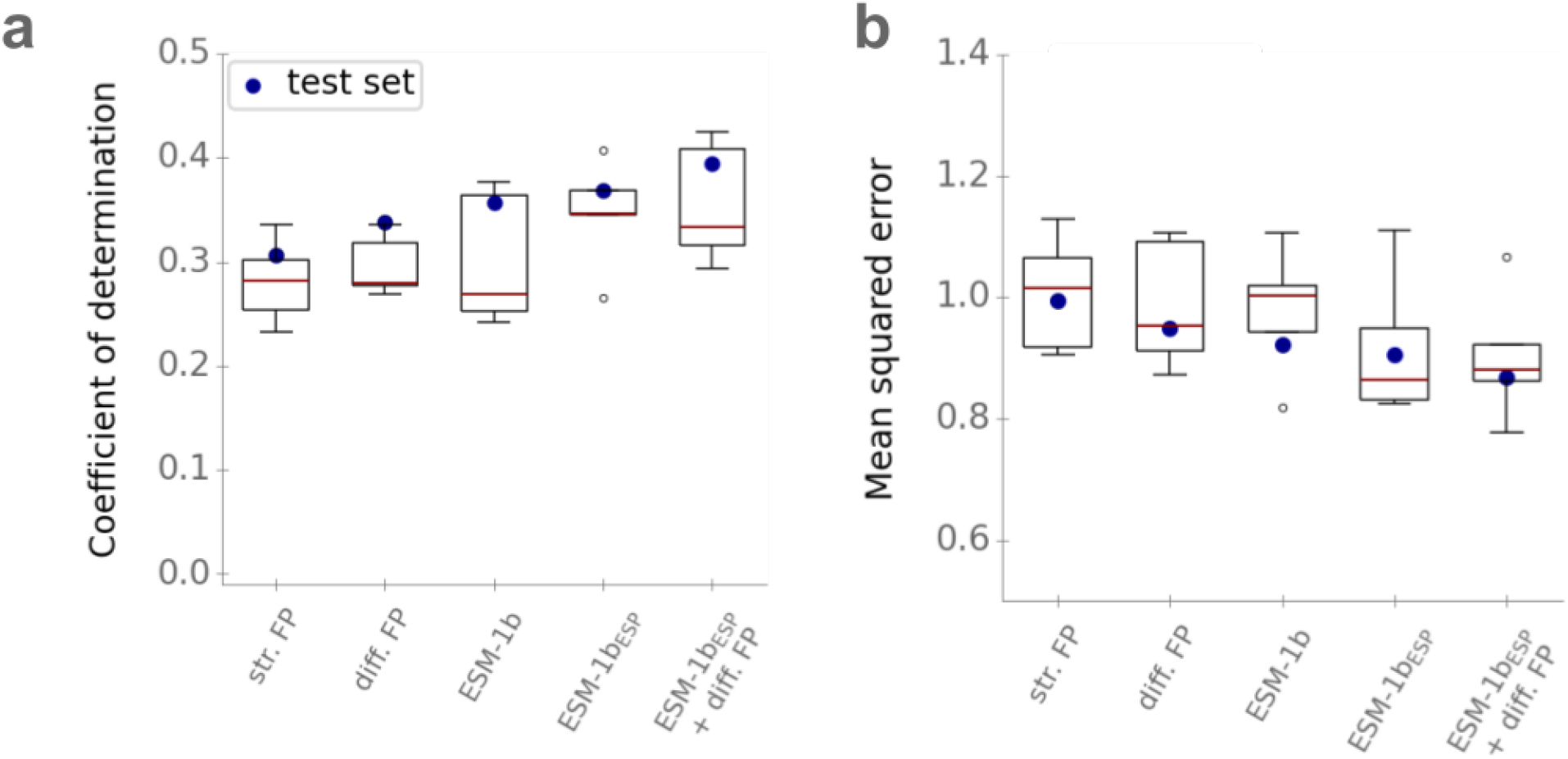
Using enzyme and reaction information combined leads to improved *k*_cat_ predictions. **(a)** Coefficients of determination *R*^2^ for models with different inputs. **(b)** Mean squared errors (*MSE*) on log10-scale. Boxplots summarize the results of the 5-fold CVs on the training set with the best set of hyperparameters; blue dots show the results on the test set using the optimized models trained on the whole training set. Model performances are plotted for the models with structural reaction fingerprints (str. FP), difference reaction fingerprints (diff. FP), *ESM-1b* vectors (*ESM-1b*), task-specific *ESM-1b* vectors (*ESM-1bESP*), and with enzyme and reaction information (*ESM-1bESP* + diff. FP).

To test if the better performance of difference fingerprints is statistically significant, we used a two-sided Wilcoxon signed-rank test that compared the absolute errors of the two models on the test set, resulting in *p* = 0.0089. Hence, using difference reaction fingerprints leads to statistically significant improvements, and we chose the difference reaction fingerprints to represent the catalyzed chemical reactions in the further analyses.

### Numerical enzyme representations alone lead to reasonable *k*_cat_ predictions

The turnover number *k*_cat_ of an enzyme-catalyzed reaction is highly dependent on the catalyzing enzyme. It can vary by orders of magnitude even between isoenzymes that catalyze the same reaction but differ in amino acid sequence^26^. To account for this dependence when predicting *k*_cat_, it is crucial to create meaningful enzyme representations as inputs to machine learning models. In recent years, deep learning architectures that were originally developed for natural language processing (NLP) tasks, such as translating a sentence from one language into another, have been applied successfully to the creation of numerical protein representations from amino acid sequences^17,27^. When applied to natural languages, NLP models typically represent all words in a sentence through numerical vectors that encode information about the words’ contents and positions. When applying NLP models to protein sequences, proteins replace sentences and amino acids replace words.

The current state-of-the-art architecture for NLP tasks is a Transformer Network^28^, which can, in contrast to previous methods, process all words of a sequence with arbitrary length simultaneously. The Facebook AI Research team trained such a Transformer Network, called *ESM-1b*, with a dataset of ∼ 27 million protein sequences from the UniRef50 dataset^29^ to create 1280-dimensional numerical protein vectors. The *ESM-1*b model was trained in a self-supervised fashion, i.e., 10-15 % of the amino acids in a sequence were masked at random, and the model was trained to predict the identity of the masked amino acids. It has been shown that the resulting representations contain rich information about the structure and the function of the proteins^17,30,31^. Using the pre-trained *ESM-1*b model^17^, we calculated these 1280-dimensional representations for all enzymes in our dataset, in the following referred to as *ESM-1b* vectors.

In a previous project^30^, we created a fine-tuned and task-specific version of the *ESM-1b* model that led to improved predictions for the substrate scope of enzymes, a problem for which abundant training data exists. Such comprehensive data is required to re-train the *ESM-1b* model, but is not available for *k*_cat_, and we were thus unable to create a version specific to the task of predicting *k*_cat_. However, we speculated that the *ESM-1b* vectors fine-tuned previously for the prediction of enzyme-substrate pairs might also improve *k*_cat_ predictions. To test this hypothesis, we used our previously published model, ESP^30^, to calculate fine-tuned representations for all enzymes in our dataset. In the following, we will refer to these representations as *ESM-1b*_*ESP*_vectors.

We tested how well models that use enzyme information alone can predict turnover numbers. We trained a gradient boosting model^32^ that used either the *ESM-1b* or *ESM-1b*_*ESP*_ vectors to predict the *k*_cat_ value of enzyme-catalyzed reactions, without using any additional information on the reaction or on substrates or products. Gradient boosting models consist of many decision trees that are built iteratively during the training process. In the first iteration, a single decision tree is built that tries to predict the correct *k*_cat_ for all data points in the training set. In all following iterations, a new decision tree is built in order to reduce the errors that have been made by the already existing trees. After training, many different decision trees exist that ideally focus on different aspects of the input features and that try to predict the correct outcome as an ensemble^33^.

To optimize the hyperparameters of the gradient boosting models, we again performed 5-fold cross validations (CV) with a random grid search on the training set. Afterwards, we re-trained each model with its best hyperparameters on the whole training set. On the test set, the model with *ESM-1b* vectors as its input achieves a coefficient of determination *R*^2^ = 0.36, a mean squared error *MSE* = 0.92, and a Pearson correlation coefficient *r* = 0.60 (**Figure 3**). The model with *ESM-1b*_*ESP*_ vectors achieves slightly improved performance, with *R*^2^ = 0.37, *MSE* = 0.91, and *r* = 0.61 on the test set (**Figure 3**). Thus, a model based on enzyme information alone leads to slightly better *k*_cat_predictions than a model based only on information on the catalyzed chemical reaction.

We had hypothesized that task-specific *ESM-1b*_*ESP*_ vectors would lead to improved results compared to the original *ESM-1b* vectors. Indeed, the model with *ESM-1b*_*ESP*_ input vectors led to slightly improved *R*^2^, *MSE*, and Pearson r on the test set and achieved better results during CV (**Figure 3**). To test if this performance difference is statistically significant, we used a one-sided Wilcoxon signed-rank test that compared the absolute errors made by both models on the test set, resulting in p = 0.41. Although the difference in absolute errors is not statistically significant at the commonly used 5% level, p *<* 0.5 indicates that rejecting the null hypothesis (*ESM-1b*_*ESP*_ vectors do not lead to superior results) is more likely than not to improve the model. Thus, we chose to represent enzymes through *ESM-1b*_*ESP*_ vectors in the following.

### A joint model with enzyme and reaction information leads to improved *k*_cat_ predictions

To train a Turnover Number Prediction model (TurNuP) with enzyme and reaction information, we concatenated the *ESM-1b*_*ESP*_ vector and the difference reaction fingerprint for every data point in our dataset. We used this resulting vector as the input for a gradient boosting model. As before, we performed a 5-fold CV with a random grid search for hyperparameter optimization, trained the model with the best set of hyperparameters on the whole training set, and validated it on the test set. The final TurNuP model achieves a coefficient of determination *R*^2^ = 0.40, a mean squared error *MSE* = 0.86, and a Pearson correlation coefficient r = 0.63 on the test set (**Figures 3** and **4**).

**Figure 4.**
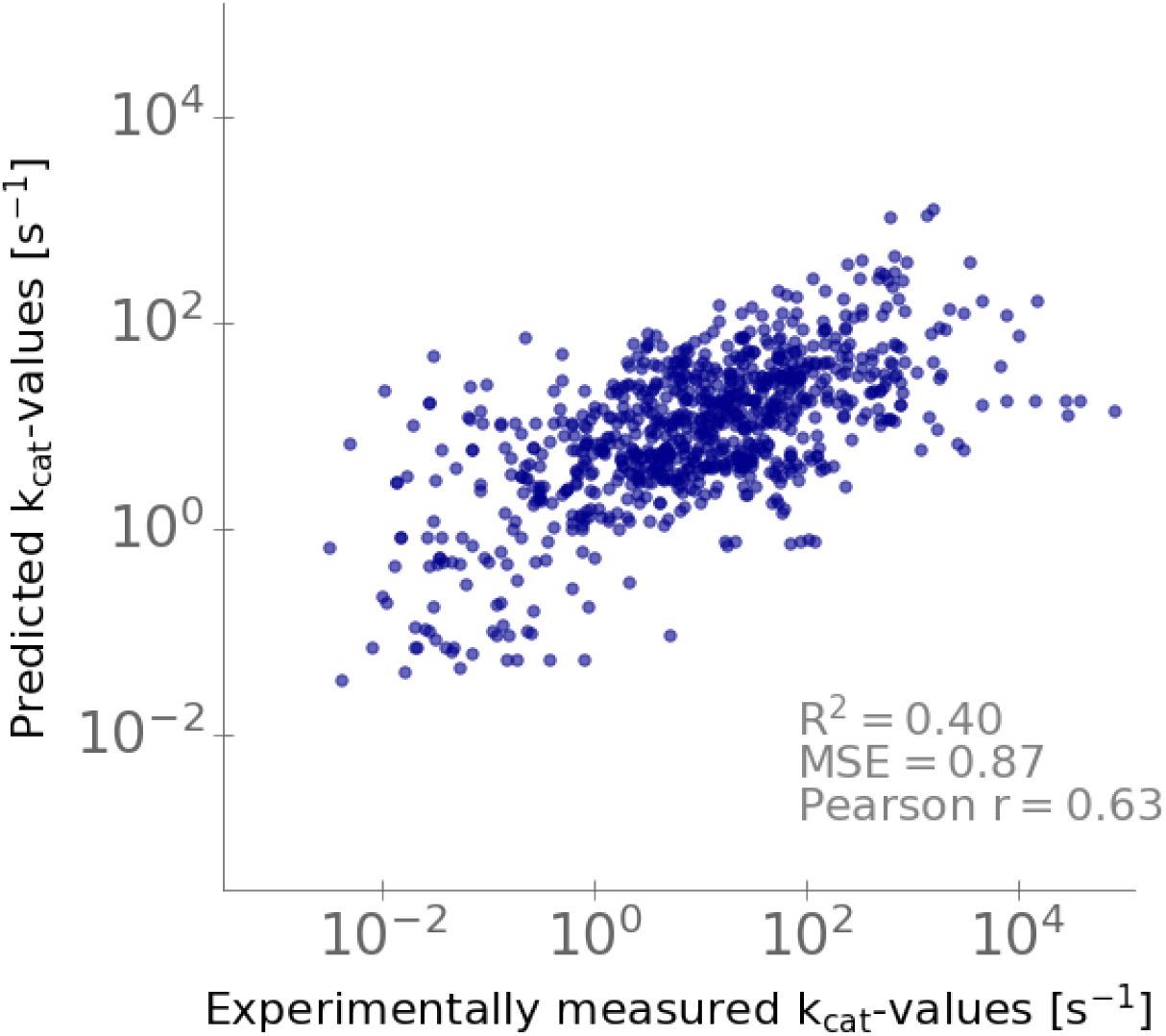
Comparison of predicted and experimentally measured *k*_cat_ values. *k*_cat_ values predicted with the complete TurNuP model, plotted against the corresponding experimental measurements. Each dot is one data point from the test set.

Using enzyme and reaction information combined in one model improves performance compared to using only enzyme or only reaction information (**Figure 3**). To compare these differences statistically, we used a one-sided Wilcoxon signed-rank test, testing if the the absolute errors on the test set for the joint model are lower than for the models with either only enzyme or only reaction information. These tests showed that the differences are statistically significant at the 5% level, with p = 0.043 (difference fingerprint) and *p* = 0.0046 (*ESM-*1b_*ESP*_). However, the improvement for the joint model is relatively small, indicating that the information stored in the reaction fingerprints and in the enzyme representations are overlapping. This overlap is not surprising, as the enzyme sequence contains information about the catalyzed reaction^30^; conversely, given that enzymes evolve on fitness landscapes shaped by the catalyzed reactions, the chemical reaction likely also contains information about the type of catalyzing enzyme.

### TurNuP provides meaningful predictions even if no close homologs with known *k*_cat_ exist

In our study on predicting the substrate scope of enzymes^30^, we found that prediction performance depends strongly on the sequence similarity between a target enzyme and enzymes in the training set, consistent with the widely held belief that enzymes are more likely to be functionally similar if they have more similar sequences [34]. We hence examined the performance of TurNuP for enzyme sets that differed in their maximal similarity to proteins in the training set. We partitioned the enzymes in the test set into four subsets with 0-40%, 40-80%, 80-99%, and 99-100% maximal sequence identity to enzymes in the test set, respectively. We calculated TurNuP’s coefficient of determination for all four categories (**Figure 5a**, black points). As expected, prediction performance decreases with increasing distance of the enzyme’s amino acid sequence to proteins in the training set. While TurNuP’s coefficient of determination is *R*^2^ = 0.66 for 99-100% sequence identity, it decreases to *R*^2^ = 0.28 for enzymes with a maximal sequence identity below 40%.

**Figure 5.**
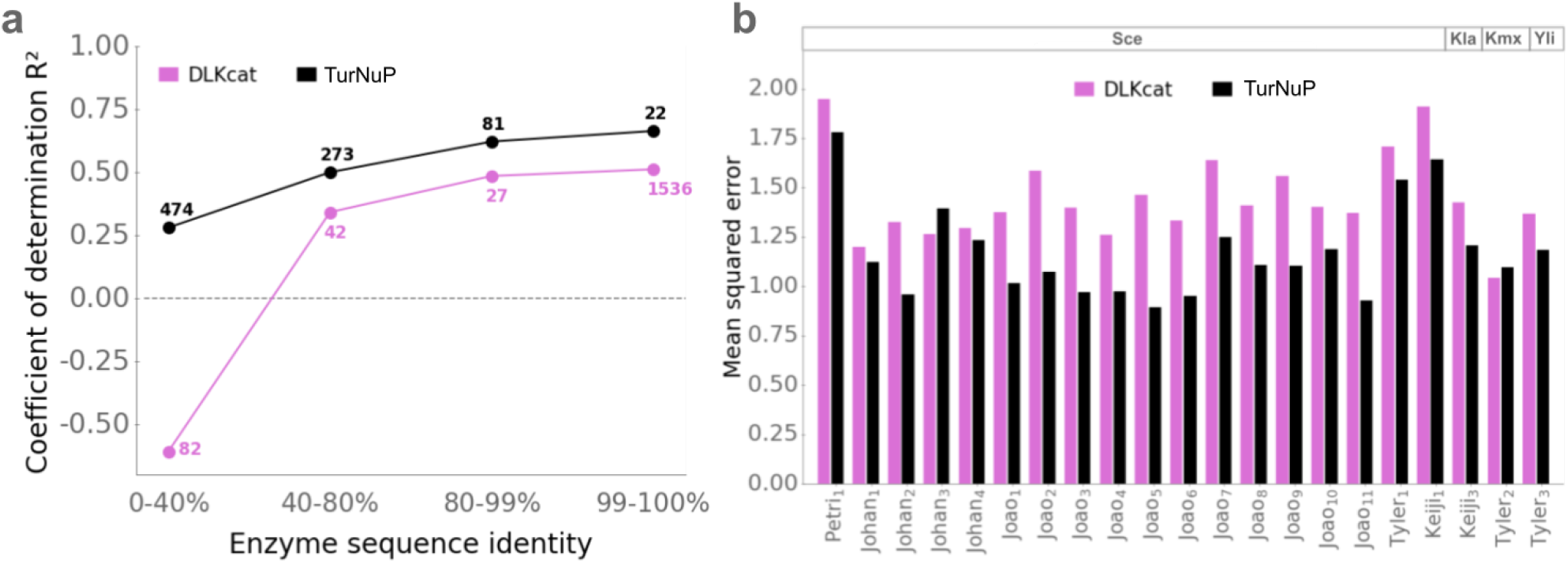
Predictions are more accurate for enzymes more similar to proteins in the training set, and TurNuP predictions are more accurate compared to an existing deep learning model. **(a)** Coefficients of determination *R*^2^ for the test sets for our TurNuP model (black) and the previously published DLKcat model^16^ (magenta) for different levels of maximal enzyme sequence identity compared to enzymes in the training set. Numbers next to points show how many data points of this category are in the test set. The horizontal dashed line corresponds to a model that predicts the same mean *k*_cat_ value for all test data points. **(b)** Mean squared errors (*MSE*) for the prediction of absolute proteome data compared to experimental data. Proteome predictions were achieved with enzyme-constrained genome-scale models, parameterized with *k*_cat_ values predicted with TurNuP (black) or with the DLKcat model (magenta). Proteome data was predicted for four different yeast species (Sce, *Saccharomyces cerevisiae*; Kla, *Kluyveromyces lactis*; Kmx, *Kluyveromyces marxianus*; Yli, *Yarrowia lipolytica*) in 21 different culture conditions (for details, see Methods).

A simple, straight-forward, and often used alternative method to predict approximate *k*_cat_ values is to simply average over the *k*_cat_ values of the most similar enzymes. Such simple averages are expected to work well in cases where kinetically characterized homologs with highly similar amino acid sequences exist; in contrast, they are unlikely to provide good estimates if no close homologs with known *k*_cat_ exist. As expected, for enzymes in the test set with close homologs in the training set (99-100% max. identity), the geometric mean across the three most similar enzymes in the test set leads to reasonable estimates, with *R*^2^ = 0.21 (*N* = 22). In contrast, averaging over the three most similar enzymes leads to a dismal *R*^2^ = 0.02 (*N* = 474) if no close homologs exist in the training data (0-40% max. identity).

These results demonstrate that any sophisticated prediction model for turnover numbers will be most relevant for enzymes for which no close homologs with known *k*_cat_ exist. As expected, TurNuP predictions are statistically significantly better than those provided by simple averages, across the complete test set (*R*^2^ = 0.40 vs. *R*^2^ = 0.24, *N* = 851, *p* = 0.00040 from one-sided Wilcoxon signed-rank test) as well as for the similarity classes (0-40% max. identity: *p* = 2.2 × 10^−7^, *N* = 22; 99-100% max. identity: *p* = 0.0054, *N* = 474).

### TurNuP outperforms previous models for predicting *k*_cat_

Heckmann et al.^14^ trained and validated a machine learning model for the prediction of *k*_cat_ values for *E. coli*. As enzyme-related input features, their model used enzyme molecular weight and global structural disorder, as well as several molecular details of the active site: number of residues, solvent access, depth, hydrophobicity, secondary structure, and exposure. Additional input features were reaction flux, number of substrates, the dissociation constant *K*M, EC number, substrate and product concentration, thermodynamic efficiency, and the pH value and temperature at which *k*_cat_ was measured *in vitro*. Out of this large set of features, the most important input was found to be the reaction flux, which was calculated by performing parsimonious flux balance analyses (pFBA)^35,36^. The total number of training and validation data points was limited to 215, as Heckmann et al.^14^ only considered reactions from *E. coli*, and as many input features are not available for most enzymes – the least widely available features were information about the enzymes’ active site, and the pH and temperature of the *in vitro* experiment. The model achieved a coefficient of determination *R*^2^ *≈* 0.34 on a test set. In comparison, our general model, which can be applied to enzymes from any organism and which does not require any enzyme information beyond the linear amino acid sequence, achieves an improved coefficient of determination of *R*^2^ = 0.40 on a test set. The DLKcat model by Li et al.^16^ examined the same problem addressed here, the prediction of *k*_cat_ values across the space of all possible enzymatic reactions. Therefore, we undertook a more in-depth comparison to DLKcat. We partitioned the enzymes in the DLKcat test set according to their maximal sequence identity to proteins in the DLKcat training set, analogous to the classification of enzymes in our own test set. **Figure 5a** shows that TurNuP (black) achieves substantially higher coefficients of determination than DLKcat (magenta) for all categories of enzyme sequence identity. We tested if the differences in model performance are statistically significant using one-sided Wilcoxon–Mann–Whitney tests; for each subset, the test compares the distributions of absolute errors of the two models. For three of the four subsets, the improvements are statistically significant, with p-values of *p* = 1.5 × 10^−10^ (0-40%), *p* = 0.0039 (40-80%), and *p* = 0.0025 (80-99%). With *p* = 0.080, the *p*-value for the forth subset (99-100%) is slightly higher than the customary significance level of 5%; this might be related to the small sample size (*N* = 22) of the TurNuP test set in this category, caused by our decision to make sure that amino acid sequences in the test set are distinct from those in the training set.

We also compared the performance of DLKcat to the simple strategy of taking the geometric mean across the *k*_cat_ values of the three most similar enzymes in the training set. Across all data points in the test set, *k*_cat_ predictions by DLKcat (*R*^2^ = 0.45, *N* = 1687) were only marginally better than those provided by simple averages (*R*^2^ = 0.44; *p* = 0.0033 from one-sided Wilcoxon signed-rank test). This surprising similarity in prediction quality between DLKcat and simple *k*_cat_ averages is likely related to the strong sequence similarities between DLKcat’s training and test sets. 68% (*N* = 1142) of the enzymes in the test set are also included in the training set (100% max. sequence identity), and a further 23% (*N* = 394) are at least 99% identical to amino acid sequences in the training set (but not 100% identical). In contrast to TurNuP, DLKcat was not challenged during its training to predict *k*_cat_ values for enzymes with dissimilar amino acid sequences^16^. As a consequence, using the geometric mean across *k*_cat_ values of similar enzymes is a major improvement in cases where no close homologs with known *k*_cat_ exist (0-40% max. identity). While simple averages achieve *R*^2^ = 0.11 in this case, DLKcat predictions lead to *R*^2^ = *−*0.61 (*p* = 0.008, two-sided Wilcoxon signed-rank test). The strongly negative coefficient of determination shows that for enzymes without kinetically characterized close homologs in the training data, DLKcat predictions are substantially worse than a trivial model that “predicts” the same mean *k*_cat_ value independent of the enzyme and the reaction. In sum, using DLKcat appears to be no improvement over the simple approach of calculating an average *k*_cat_ value across the most similar enzymes with available turnover numbers.

Although TurNuP performs better than DLKcat in each of the four categories of enzyme sequence identities (**Figure 5a**), DLKcat achieves a higher *R*^2^ value (*R*^2^ = 0.44) on its overall test set, compared to the TurNuP model on its overall test set (*R*^2^ = 0.40). This counter-intuitive observation is an example of Simpson’s paradox. It is caused by the differential distribution of data points across categories in the DLKcat and TurNuP test sets. As shown by the numbers above/below points in **Figure 5a**, 91% of the data points in the DLKcat test set fall into the 99-100% identity class, while the majority of data points in the TurNuP test set (56%) have less than 40% sequence identity to any enzymes in the corresponding training set and are hence much harder to predict.

Li et al.^16^ designed a pipeline to use predicted *k*_cat_ values for the parameterization of enzyme-constrained genome-scale metabolic models, with the goal of predicting the proteome allocation patterns of yeast species. They compared the resulting proteome predictions to absolute proteomics measurements for four yeast species in 21 different environments. We employed this pipeline to test if *k*_cat_ values calculated with the TurNuP model lead to improved proteome predictions. In 19 out of 21 environment-species combinations, our *k*_cat_ values led to improved predictions (*p* = 0.00010, one-sided binomial test). The mean squared errors between measured and predicted protein abundances improved on average by ∼ 18% when using TurNuP (**Figure 5b**).

### Using additional input features does not improve model performance

TurNuP employs very general input features, using only the enzyme’s linear amino acid sequence and information on the reaction’s substrates and products. However, it is unclear if these features cover all important aspects for predicting *k*_cat_. To test if we can improve prediction quality, we examined three potential additional input features: Michaelis constants *K*_M_, Codon Adaptation Indices (CAIs), and reaction fluxes.

The Michaelis constant *K*_M_ is defined as the substrate concentration at which an enzyme works at half of its maximal catalytic rate; hence, *K*_M_ quantifies the affinity of an enzyme for its substrate. It has been shown that *k*_cat_ is correlated with the enzyme’s Michaelis constant(s) of the reaction’s substrate(s)^23^. To utilize this correlation for the prediction of *k*_cat_, we determined *K*_M_values for all enzyme-substrate combinations in our dataset. Where available, we extracted suitable *K*_M_values from the BRENDA database^20^ (∼ 7% of the enzyme-substrate pairs in our dataset); for all other cases we applied a machine learning model that uses numerical representations of the substrate and the enzyme as its input to predict *K*_M_^37^. For reactions with multiple substrates, we took the geometric mean of all *K*_M_ values to obtain a single *K*_M_ value for every data point. To calculate how much variance of *k*_cat_ can be explained by *K*_M_, we fitted a linear regression model to the training set, with the log_10_-transformed *K*_M_ value as the only input. The linear regression model achieves a coefficient of determination *R*^2^ = 0.11, a mean squared error *MSE* = 1.28, and a Pearson correlation coefficient *r* = 0.34 on the test set (**Figure S2**). We thus considered *K*M a promising candidate for improving the TurNuP predictions.

The second additional input feature, the Codon Adaptation Index (CAI), quantifies the synonymous codon usage bias of protein-coding genes. It is widely used as an indicator of gene expression and protein levels, with highly expressed genes typically using more ‘preferred’ codons than less highly expressed genes^38^. The CAI is a value between 0 and 1 that describes the similarity of synonymous codons frequencies between a given gene and a set of highly expressed genes, where values close to 1 indicate nearly optimal codon usage, typically associated with a high expression level in evolutionarily relevant environments. We calculated CAI for all enzymes in our dataset originating from *E. coli*. We fitted a linear regression model to the corresponding 237 data points in the training set, with CAI as the only input feature. We validated the model on 66 test data points (**Figure S3**). The model achieved a coefficient of determination *R*^2^ = 0.012, a *MSE* = 1.31, and a Pearson correlation coefficient *r* = 0.12 on the test set, indicating that CAI cannot explain much of the variance of *k*_cat_ values. Hence, we did not consider CAI a promising candidate for improving the TurNuP predictions, and we did not calculate CAI for other organisms beyond *E. coli*.

The most important input feature in the *k*_cat_ prediction model established by Heckmann et al. for reactions in *E. coli*^14^ was an estimate of the reaction flux, calculated using parsimonious flux balance analysis (pFBA)^35,36^ across a broad range of nutrient conditions. For 108 metabolic genome scale models from the BiGG database^39^, we calculated fluxes in a similar way as Heckmann et al. (Methods). For further analyses, we selected the six BiGG models of distinct species that showed the highest Pearson correlation between predicted fluxes and measured *k*_cat_ values in the training set. We mapped the calculated fluxes to values from our dataset. In cases where no metabolic genome scale model was available for an organism, we mapped the flux of an identical reaction but from a different organism to the data point. If we were not able to find the identical reaction in the BiGG database, we selected the most similar one using a similarity score (see Methods). To calculate how much variance of the *k*_cat_ values can be explained by the calculated fluxes, we fitted a linear regression model to the training set, with the log *k*^10^-transformed fluxes as the only input. The fitted model achieves a coefficient of determination *R*^2^ = 0.021, a *MSE* = 1.40, and a Pearson correlation coefficient *r* = 0.15 on the test set (**Figure S4**). Thus, we found no evidence for a high predictive power of fluxes beyond *E. coli*; however, as fluxes were the most important predictor for *k*_cat_ in Ref.^14^, we still retained them as a potential additional input feature for TurNuP.

To test if adding *K*_M_ and reaction flux as input features improves model performance, we trained a new model. As the model input, we created a concatenated vector comprised of the enzyme *ESM-1b*^*ESP*^ vector, the difference reaction fingerprint, the reaction flux, and the Michaelis constant *K*_M_ for every data point. For a gradient boosting model, we then performed a 5-fold CV with a random grid search for hyperparameter optimization. Afterwards, we trained the model with the best set of hyperparameters on the complete training set. On the test set, this model achieves a coefficient of determination *R*^2^ = 0.39, a *MSE* = 0.87, and a Pearson correlation coefficient *r* = 0.63. Thus, model performance did not improve compared to the model without the additional input features flux and *K*_M_.

### The TurNuP web server provides an easy acces to the prediction model

We implemented a web server that facilitates an easy use of the TurNuP model without requiring programming skills or the installation of specialized software. It is available at https://turnup.cs.hhu.de. As input, the web server requires an enzyme amino acid sequence and representations of all substrates and all products; the latter can be provided either as SMILES strings, KEGG Compound IDs, or InChI strings. Users can enter a single enzymatic reaction into an online form, or upload a CSV file with multiple reactions. Since TurNuP was trained only with natural reactions of wild-type enzymes, we recommend to use the web server only for such enzyme-reaction pairs.

## Discussion

Predicting the turnover number of enzyme-catalyzed reactions is a complex task, and the available datasets for model training are small and noisy. For example, Bar-Even et al.^23^ found that that up to 20% of the entries in BRENDA differ from the entries in the reference papers, probably caused by copying errors and erroneous replacements of units. Even aside from such obvious errors, the variance of *k*_cat_ measurements for the same enzyme-reaction pairs between different studies can be high. We found an average deviation of 5.7-fold (mean deviation on log_10_-scale = 0.75) between two *k*_cat_ measurements for the same enzyme-reaction pair. This variance is likely not only due to errors in the databases, but also to different experimental procedures or varying assay conditions, such as temperature and pH value. When comparing a single measurement to the geometric mean of all other measurements for the same enzyme-reaction pair, we found an average deviation of 3.3-fold (mean deviation on log10-scale = 0.52). This compares to an average deviation of 5.1-fold (mean deviation on log10-scale = 0.71) of predicted *k*_cat_ values with our TurNuP model compared to the geometric mean of all available measurements for this enzyme-reaction pair. These numbers indicate that in practice, using predictions calculated with the TurNuP model may lead to similar deviations and error rates compared to performing experimental measurements.

Although the accuracy of TurNuP’s predictions is not very different from that of experimental estimates, model accuracy can still be improved. On the one hand, we trained and validated the model with a total of only 4 271 data points, which is rather small for a machine learning model with high-dimensional input vectors. Once more high-quality training data becomes available, model performance will most likely improve. On the other hand, *k*_cat_ values can differ widely if measured under different experimental conditions such as varying pH and temperature. However, as information about the experimental conditions is mostly unavailable in databases for enzyme kinetic parameters, we were not able to include these conditions as an input to our prediction model. Manually extracting this information from research papers has the potential to further improve accuracy of the prediction models and to create models that are capable of accounting for different experimental conditions. Moreover, as we have shown above, the high variability of experimental estimates for the same enzyme-reaction pairs indicates a lot of noise across the measured *k*_cat_ values. Better predictions will become possible in the future if experimental variation will be reduced through improved technologies.

TurNuP achieves superior performance compared to previous methods for predicting *k*_cat_. Its coefficient of determination (*R*^2^ = 0.4) is higher than than of Heckmann et al. (*R*^2^ *≈* 0.34)^14^, who trained an organism-specific prediction model with very detailed and expert-crafted input features, including enzyme active site properties, metabolite concentrations, reaction fluxes, and experimental conditions. TurNuP also outperforms the most recent method for predicting *k*_cat_, the DLKcat model^16^ (**Figure 5**). One reasons for TurNuP’s superior performance might be the use of state-of-the-art enzyme representations compared to convolutional neural networks (CNNs) and the use of representations for the whole chemical reactions instead of using only information on one of the substrates.

An additional important reason might be a careful preprocessing of the *k*_cat_ dataset. We excluded all data points with mutated enzymes and non-natural reactions, because we are mainly interested in predicting turnover numbers for natural reactions of wild-type enzymes, and we hypothesized that we do not have enough training data to teach our model to predict the catalytic effect of enzyme mutations or to predict the *k*_cat_ value of non-natural enzyme-reaction pairs. When including mutated enzymes, one has to be very careful when splitting the dataset into training and test set, as one may easily end up with many nearly identical enzymes in both sets. 91% of the enzymes in the DLKcat test set have a maximal sequence identity between 99 and 100% compared to the enzymes in the training set. It is likely that the same issue arose in the validation set used for hyperparameter optimization; such a structure of training and validation sets makes it difficult to train a model that generalizes well to enzymes not highly similar to those in the training set. Indeed, we showed that DLKcat does not produce meaningful predictions for enzymes with a maximal sequence identity lower than 40% compared to the enzymes in the training set, and total model performance on the test set is not meaningfully better than calculating *k*_cat_ averages across the most similar enzymes in the training set. In comparison, less than 3% of the enzymes in the TurNuP test set have amino acid sequences that are *>*99% identical to those of homologs in the training data. Moreover, TurNuP generalizes well even to enzymes that are not highly similar to enzymes in the training set, and provides a major improvement over simple *k*_cat_ averages.

To achieve these results, we used general input features: the *ESM-1b*_*ESP*_ vector^17^, a fine-tuned, state-of-the-art numerical representation of the enzyme, calculated from its amino acid sequence; and a reaction fingerprint that integrates structural information about all substrates and products^24^, which allowed us to create input vectors of fixed length even for varying numbers of reactants. Surprisingly, we found that the *ESM-1b*_*ESP*_ vectors work very well if used as the only input feature (**Figure 3**). The reason for that is neither that enzymes with high fluxes or with high expression levels have different enzyme representations, because these features by themselves do not predict much variance of the *k*_cat_ values (see Results, “Using additional input features does not improve model performance”).

It is at first sight surprising that the reaction fluxes estimated with pFBA do not explain much of the variance of *k*_cat_, while they were found to be the best predictor in the model developed by Heckmann et al.^14^. When calculating genome-scale reaction fluxes for different organisms, for many data points, we obtained fluxes that were zero or close to zero. In contrast, Heckmann et al. focused on a small dataset that mostly consisted of well-studied, central reactions in *E. coli*. Those reactions typically have fluxes substantially different from zero at least in some of the simulated conditions. It appears likely that this biased construction of a small dataset in Ref.^14^ is responsible for the high correlation observed between reaction fluxes and *k*_cat_ by Heckmann et al..

Computational estimates of *k*_cat_ values are highly relevant for the functional and kinetic study of individual enzymes^40^, and TurNuP can provide a first estimate of *k*_cat_ before performing labor-intensive experiments. Another major use case of TurNuP is the prediction of *k*_cat_ values for genome-scale metabolic models. We found that our predictions can be used successfully to improve proteome allocation predictions (**Figure 5b**). In future work, *k*_cat_ predictions with TurNuP can be combined with an existing approach for predicting Michaelis constants (*K*_M_)^37^. This would facilitate full parameterizations of non-linear enzyme kinetics in genome-scale metabolic models, a powerful tool for gaining fundamental insights into cellular physiology^9,41^.

## Methods

### Software and code availability

All software was coded in Python^42^. We created the enzyme representations using the deep learning library PyTorch^43^. We fitted the gradient boosting models using the library XGBoost^32^. We used the web framework Django^44^ to implement the TurNuP web server. The code used to generate the results of this paper, in the form of Jupyter notebooks, as well as all datasets, are available from https://github.com/AlexanderKroll/Kcat_prediction.

### Downloading *k*_cat_ data

We used data from three different databases, Sabio-RK, UniProt, and BRENDA, to create a *k*_cat_ dataset for model training and validation. We downloaded 3 971 *k*_cat_ values for wild-type enzymes together with UniProt IDs and reaction information from Sabio-RK^22^. We tried to map all metabolites involved in the reactions to unique identifiers using either a KEGG reaction ID^45^, if available, or using the metabolite names and the PubChem synonym database^46^. We removed all data points for which we could not map all substrates and all products to an ID. This resulted in a dataset with 2 830 data points for 289 different enzymes.

We downloaded 5 664 *k*_cat_ values for wild-type enzymes together with UniProt IDs and CHEBI reaction IDs from UniProt via the UniProt mapping service^21^. We mapped the metabolites of all reactions to unique IDs using CHEBI reaction IDs^47^. We removed data points, if we could not map all metabolites of a reaction to an ID. This resulted in a dataset with 1 738 *k*_cat_ values for 1 017 different enzymes.

We downloaded 14 165 turnover numbers for wild-type enzymes with protein information and substrate names from BRENDA^20^. Most of the *k*_cat_ values in BRENDA are not assigned with a unique reaction equation and the entered *k*_cat_ values are known to be prone to errors^23^. To overcome these issues, we manually checked for more than half of all points if the stated *k*_cat_ value is identical to the value from the original paper and we assigned a unique reaction equation to all manually checked data points. After removing those data points with incomplete reaction information and non unique enzyme IDs, 8 267 data points were left for 3 149 different enzymes.

### Data preprocessing

We merged all three *k*_cat_ datasets from BRENDA, Sabio-RK, and UniProt, which resulted in a dataset with 12 835 data points. We removed 1 050 duplicated data points from this data set. To obtain protein sequences for all enzymes, we used the UniProt mapping service^21^ to map all UniProt IDs to amino acid sequences. We used the Python package Bioservices^48^ to map all metabolites to InChI strings^49^. If multiple *k*_cat_ values existed for the same enzyme-reaction combination, we took the geometric mean across these values. For the calculation of the geometric mean, we wanted to ignore those values that were likely obtained under non-optimal conditions. Thus, we excluded *k*_cat_ values smaller than 1% compared to the maximal *k*_cat_ value for the same enzyme-reaction combination. Calculating the geometric mean resulted in a dataset with 7 496 entries.

The BRENDA, UniProt, and Sabio-RK databases contain many *k*_cat_ values that were measured for secondary, non-natural reactions of enzymes. As we are only interested in measurements for the natural reaction of an enzyme, we excluded *k*_cat_ values if another measurement existed for the same enzyme but for a different reaction with a *k*_cat_ value that was more than ten times higher. To further exclude data points that were measured under non-optimal conditions or for non-natural reactions of the enzyme, we excluded data points if we could find a measurement for the same reaction or the same EC number that was more than 100 times higher. The described procedures led to the removal of 3 092 data points.

We calculated reaction fingerprints and enzyme representations for all enzyme-reaction pairs (see below) and removed all 26 data points, where either the reaction fingerprint or the enzyme representation could not be calculated.

To exclude data points with possibly wrongly assigned reaction equations, we removed those 52 data points where the sum of molecular weights of substrates did not match the sum of molecular weights of the products. We removed another 55 data points because their *k*_cat_ values are outliers (i.e., values below 10^−2.5^/*s* or higher than 10^5^/*s*). This resulted in a final dataset with 4 271 data points.

### Splitting the dataset into training and test set

We randomly split the dataset into 80% training data and 20% test data. We made sure that the same enzyme would not occur in the training and the test set. We further split the training set into 5 disjoint subsets for a 5-fold cross-validation (CV) to perform hyperparameter optimizations of the machine learning models. In order to achieve a model that generalizes well during CV, we created these 5 subsets also in such a way that the same enzyme did not occur in two different subsets.

### Calculating enzyme representations

To create the *ESM-1b* model^17^, the Facebook AI research (FAIR) team trained a Transformer Network^28^ with 33 hidden layers and a hidden layer size of 1 280 using ∼ 27 million protein amino acid sequences from the UniRef50 dataset^29^. To process a protein sequence, the type and position of every amino acid in a sequence is encoded in a 1 280-dimensional numerical vector. All amino acid representations of a sequence are simultaneously applied to the *ESM-1b* model and updated for 33 time steps using the attention mechanism^28^. The attention mechanism allows to use all representations as an input when updating a single amino acid representation. The attention mechanism selectively chooses only relevant input when calculating an update of a representation. To train the *ESM-1b* model, randomly 10 *−* 15% of the amino acids in a sequence are masked. The model is then trained to predict the type of the masked amino acids. After training, a single representation for the whole model can be created by calculating the element-wise mean of all amino acid representations after they were updated for 33 times.

We used the trained *ESM-1b* model and the code provided on the GitHub repository of the FAIR team^17^, to calculate a 1 280-dimensional numerical representation for every enzyme in our dataset. As the *ESM-1b* model can only process amino acid sequences up to 1 024 amino acids, we only used the first 1 024 amino acids for those sequences that were too long.

To calculate the fine-tuned enzyme representations that were originally created for the task of predicting the substrate scope of enzymes^30^, the *ESM-1bESP* vectors, we used code and models provided on the following GitHub repository: https://github.com/AlexanderKroll/ESP.

### Calculating reaction fingerprints

To calculate difference and structural reaction fingerprints, we first represented all reactions in our dataset using the language SMARTS^50^. SMARTS can be used to describe patterns of small molecules and of chemical reactions. To calculate the reaction fingerprints, we used functions from the RDKit^24^ package Chem with the reaction SMARTS as the input.

Structural reaction fingerprints are created by first calculating 1638-dimensional binary molecular fingerprints (ExplicitBitVect) for all substrates and products. Then, the bitwise OR-function is separately applied to all substrate fingerprints and to all product fingerprints, which results in two 1638-dimensional binary vectors with information about the substrates and about the products, respectively. Finally, both vectors are concatenated, which results in a 3276-dimensional binary vector with structural information about the reaction. We used the RDKit function *Chem*.*rdChemReactions*.*CreateStructuralFingerprintForReaction* to calculate the fingerprints. To calculate difference reaction fingerprints, first, a 2048-dimensional binary atom-pair fingerprint (AtompairFP) for each substrate and each product is calculated. Then, the fingerprints for all substrates and also for all products are element-wise summed. The resulting fingerprint for the products is then subtracted from the fingerprint for the substrates, which results in a 2048-dimensional reaction fingerprints with positive and negative integers. To calculate these fingerprints, we used the RDKit function *Chem*.*rdChemReactions*.*CreateDifferenceFingerprintForReaction*.

### Hyperparameter optimization for gradient boosting models

To perform hyperparameter optimizations for all gradient boosting models, we split the training set into five disjoint subsets with approximately equal sizes to perform 5-fold cross-validations (CVs). We performed a random grid search for the hyperparameters learning rate, regularization coefficients *α* and *λ*, maximal tree depth, maximum delta step, number of training iterations, and minimum child weight using the Python package hyperopt^51^. Afterwards, we chose the set of hyperparameters that led to the highest mean coefficient of determination *R*^2^ during CV.

### Comparison of *k*_cat_ predictions between the DLKcat model and TurNuP

We used code provided on a GitHub repository by Li et al.^16^ to reproduce the DLKcat model and to make predictions for their test set. We divided both, the DLKcat test set and our test set, into four different subsets according to the protein sequence identity compared to the amino acid sequences in the training sets. To achieve this, we calculated for every test sequence the maximal pairwise sequence identity compared to all sequences in the training set using the Needleman-Wunsch algorithm from the software package EMBOSS^52^. We used the coefficient of determination *R*^2^ to compare the results of TurNuP with the results of the DLKcat model.

### Predicting protein abundances using predicted *k*_cat_ values

Li et al.^16^ developed a Bayesian pipeline to use predicted *k*_cat_ values for enzyme-constrained genome-scale metabolic models to predict the proteome of yeast species. We used Matlab code provided on a GitHub repository by Li et al.^16^ to follow the same pipeline for *k*_cat_ values predicted with TurNuP. The predicted proteome allocations were compared to measured proteome data for four different species in 21 different cultural conditions. The measured proteome data was taken from six different publications^53–58^.

### Statistical tests for model comparison

To test if the difference in model performance between the TurNuP model with enzyme and reaction information compared to the models with either only enzyme or reaction information is statistically significant, we applied a one-sided Wilcoxon signed-rank test implemented in the Python package SciPy^59^. We tested the null hypothesis that the median of the absolute errors on the test set for predictions made with TurNuP, ē_1_, is greater or equal to the corresponding median for predictions made with a model with only reaction or only enzyme information, ē_2_ (*H*_0_ : ē_1_ ≥ ē_2_ vs. *H*_1_ : ē_2_ > ē_1_). We could reject *H*_0_ (*p* = 0.0003 (structural fingerprint), *p* = 0.0427 (difference fingerprint), *p* = 0.0046 (*ESM-1bESP*)), accepting the alternative hypothesis *H*_1_.

We also tested if the differences in model performance between TurNuP and the DLKcat model are statistically significant for all subsets of the test set with different enzyme sequence identity levels. We used the non-parametric one-sided Wilcoxon–Mann–Whitney test implemented in the Python package SciPy^59^ to test the null hypothesis that the prediction errors for the two models are equally distributed.

We could reject the null hypothesis for three subsets at the 5% level with p-values of *p* = 1.5 × 10^−10^(0 − 40%), *p* = 0.0039 (40 − 80%), and *p* = 0.0025 (80 − 99%), while the p-value for the forth subset (99 − 100%) was slightly above the 5% level (*p* = 0.080).

### Calculating reaction fluxes

We calculated reaction fluxes for all 108 genome-scale metabolic models (GEMs) from the BiGG database^39^. We selected those six GEMs for different organisms that showed the highest correlation between calculated fluxes through parsimonious flux balance analyses (pFBA) and *k*_cat_ values in our dataset. We selected the following six models: iECO111 1330 (*Escherichia coli*), iEK1008 (*Mycobacterium tuberculosis*), iHN637 (*Clostridium ljungdahlii*), iIT341 (*Helicobacter pylori*), iSbBS512 1146 (*Shigella boydii*), and iJN1463 (*Pseudomonas putida*).

We calculated the reaction fluxes similar to the approach by Heckmann et al.^14^ for *E. coli*. For each of the six GEMs, we simulated 10 000 minimal growth sustaining environments through pFBA^36^ using the Python package COBRApy^60^. Afterwards, we calculated for every reaction the mean of all non-zero fluxes among all simulations. In all of the 10 000 simulations, first a growth sustaining environments was created with a growth rate higher than 0.1 *h*^*−*1^ and oxygen uptake was allowed with a probability of 50% for aerobic organisms. To convert the medium into a minimal media, each metabolite of the medium was removed if growth was sustained without it. If we could not obtain a non-zero flux for a reaction in all simulations, we repeated the described procedure with a flux variability analysis (FVA)^61^ instead of a pFBA. If we could not obtain a non-zero-flux for a reaction either via pFBA or via FVA, we replaced the reaction flux with the mean of all non-zero fluxes. Python code for calculating the fluxes is available on the following GitHub repository: https://github.com/Nina181/kcat_flux_relationship.

### Mapping data points to BiGG reaction IDs

We created a list with reactions from six different metabolic genome-scale models from the BiGG database^39^ (iECO111 1330, iEK1008, iHN637, iIT341, iSbBS512 1146, iJN1463). To create this list, we downloaded a json-files for each model and we extracted all substrate names and IDs (MetaNetX or KEGG), product names and IDs, and BiGG reaction IDs. We discarded all reactions with an incomplete list of substrate or product IDs. If only a MetaNetX ID and no KEGG ID was available for a metabolite, we downloaded an InChI string^49^ for the metabolite using the MetaNetX database^62^. Next, we calculated structural reaction fingerprints for all extracted BiGG reactions using the KEGG IDs and InChI strings of the substrates and products (for details see above, “Calculating reaction fingerprints”).

To map data points from our data set to BiGG reactions IDs, we calculated a pairwise similarity score between all reactions in our dataset and all reactions from the 6 extracted BiGG models. To calculate the similarity score, we used the Python function *TanimotoSimilarity* from the RDKit package DataStructs^24^ with structural reaction fingerprints as the input. This resulted in a similarity score between 0 (no similarity) and 1 (very high similarity) for all pairs of reactions. We mapped every data point in our dataset to the BiGG reaction with the highest similarity score.

### Calculating Michaelis constants

To calculate the Michaelis constants *K*M for all enzyme catalyzed reactions in our dataset, we created a list with all enzyme-substrate pairs. We used the BRENDA database^20^ to map enzyme-substrate pairs to *K*M values via the enzymes’ amino acid sequences and via a molecular fingerprint of the substrate, called ECFP vector^63^. We were able to map a *K*M value to ∼ 7% of 8 984 enzyme-substrate pairs.

If we could not find a value for an enzyme-substrate pair in the BRENDA database, we predicted *K*M using a machine learning model^37^. The *K*M prediction model uses a graph neural network (GNN)^64,65^ to create a 50-dimensional task-specific fingerprint of the substrate. These fingerprints are used together with a 1900-dimensioanl enzyme representation, called UniRep vector^27^, as the input for a gradient boosted decision tree model^32^ to predict the *K*M value for an enzyme-substrate pair. For reactions with multiple substrates, we took the geometric mean of *K*M values to create a single *K*M value for every data point.

### Calculating the Codon Adaptation Index

The codon adaptation index (CAI) for *E. coli* was calculated according to the original definition^66^, considering ribosomal protein genes as the highly expressed genes. The sequences of ribosomal protein genes were retrieved from genome annotation of *E. coli* (NC_000913.3 from RefSeq^67^).

## Acknowledgements

We thank David Heckmann for helpful discussions. Computational support and infrastructure was provided by the “Centre for Information and Media Technology” (ZIM) at the University of Düsseldorf (Germany). This work was funded by the Deutsche Forschungsgemeinschaft (DFG,German Research Foundation) through CRC 1310, and, under Germany’s Excellence Strategy, through grant EXC 2048/1 (Project ID: 390686111).

## Author contributions

AK designed the dataset and models and performed all other analyses. NAL implemented the calculation of the reaction fluxes. XPH calculated the Codon Adaptation Index (CAI) for genes from *E. coli*. MJL conceived of and supervised the study, and acquired funding. AK wrote the initial manuscript, which was edited by AK and MJL.

## Supporting Figures S1-S4

**Figure S1.**
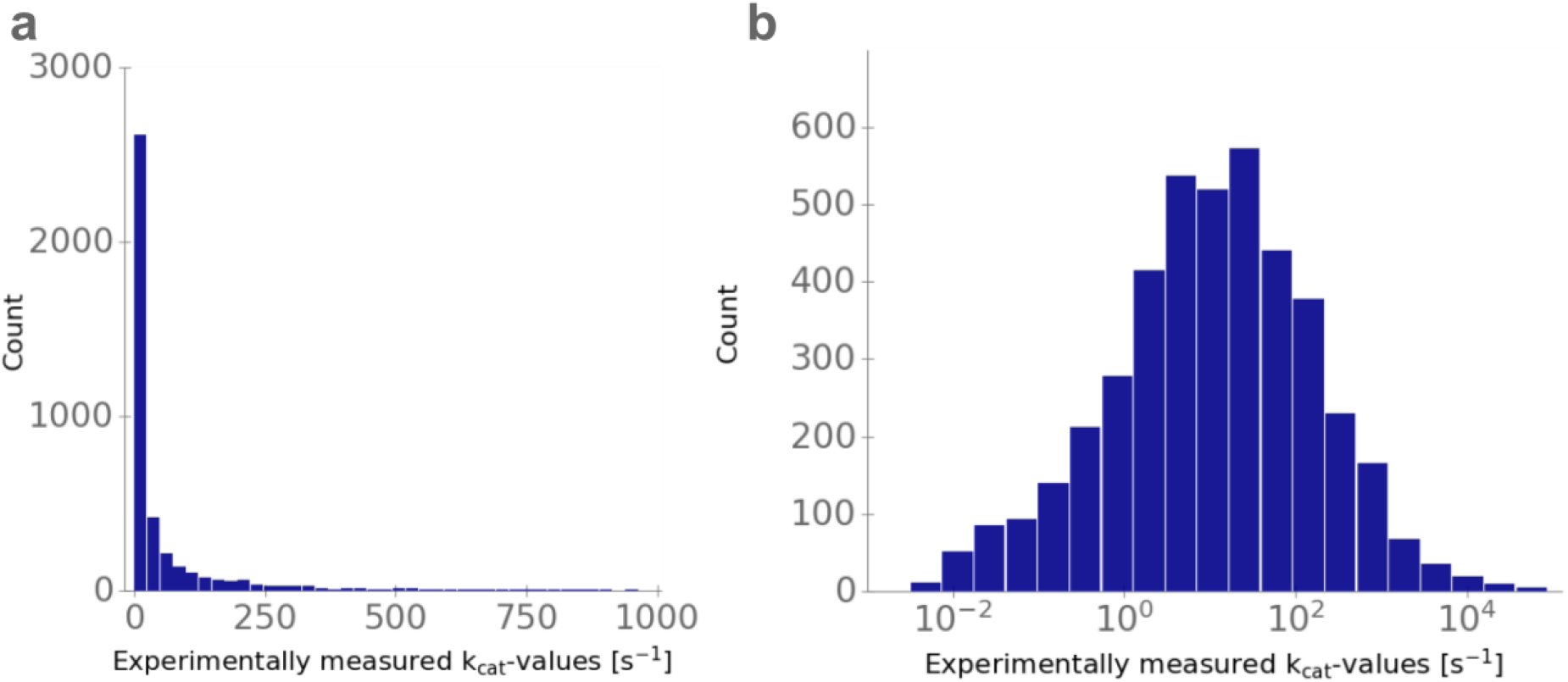
Turnover numbers are approximately log-normally distributed. Histograms of **a)** untransformed *k*_cat_ values and **(b)** *log*10-transformed *k*_cat_ values.

**Figure S2.**
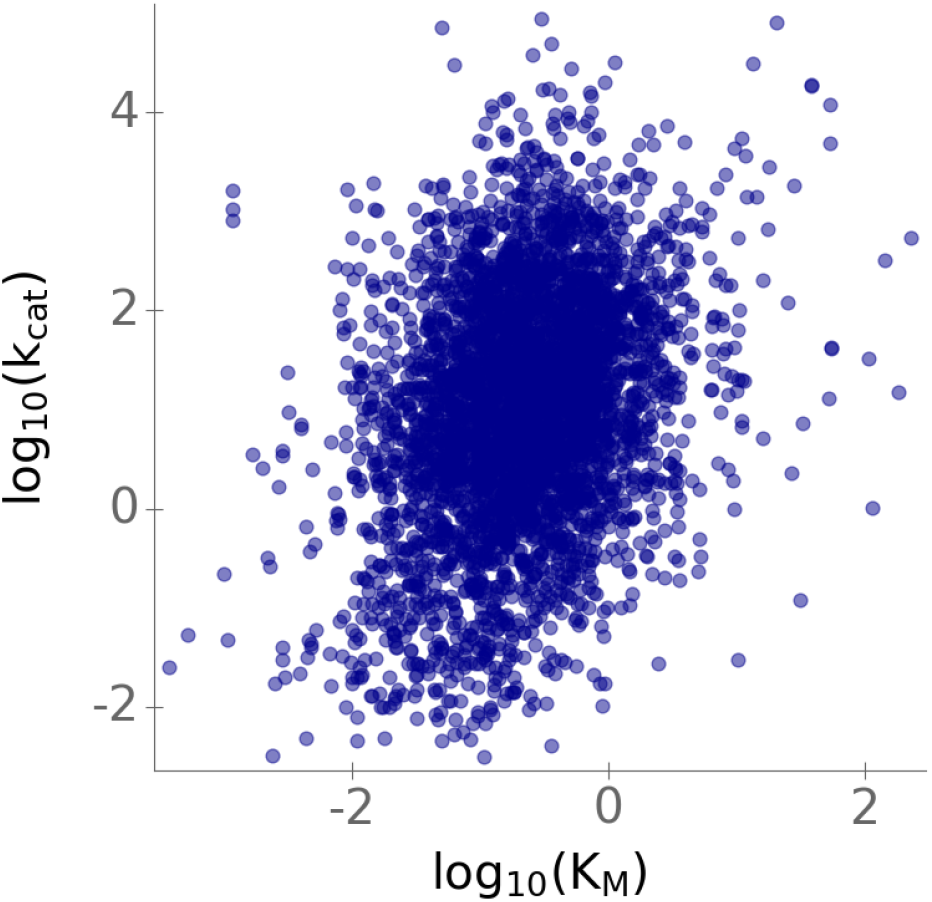
Michaelis constants *K*M compared to *k*_cat_ values. We obtained either experimentally measured *K*M values from BRENDA or predicted *K*M values for every reaction and we plotted these values against the corresponding *k*_cat_ values on a log_10_-scale. The plot contains 4271 data points from our training and test set.

**Figure S3.**
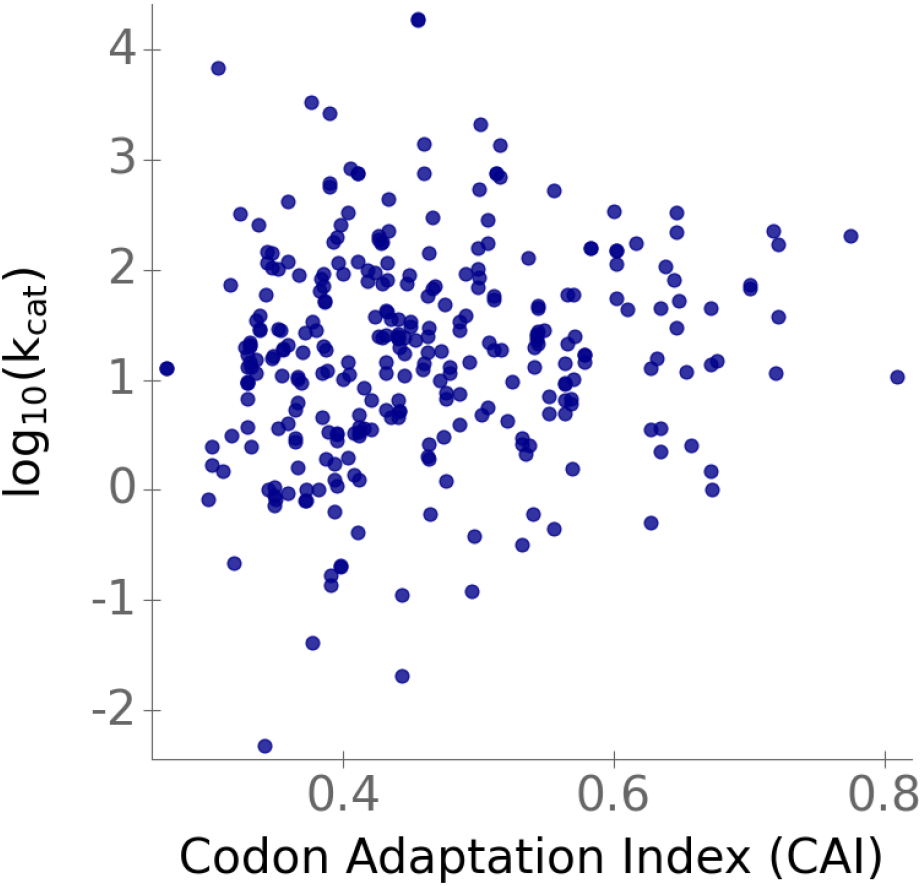
Codon Adaptation Index (CAI) for enzymes from *E. coli* compared to their *k*_cat_ values. We calculated the CAI for genes from *E. coli* and plotted it against their log_10_-transformed *k*_cat_ value. The plot contains 303 data points from our training and test set.

**Figure S4.**
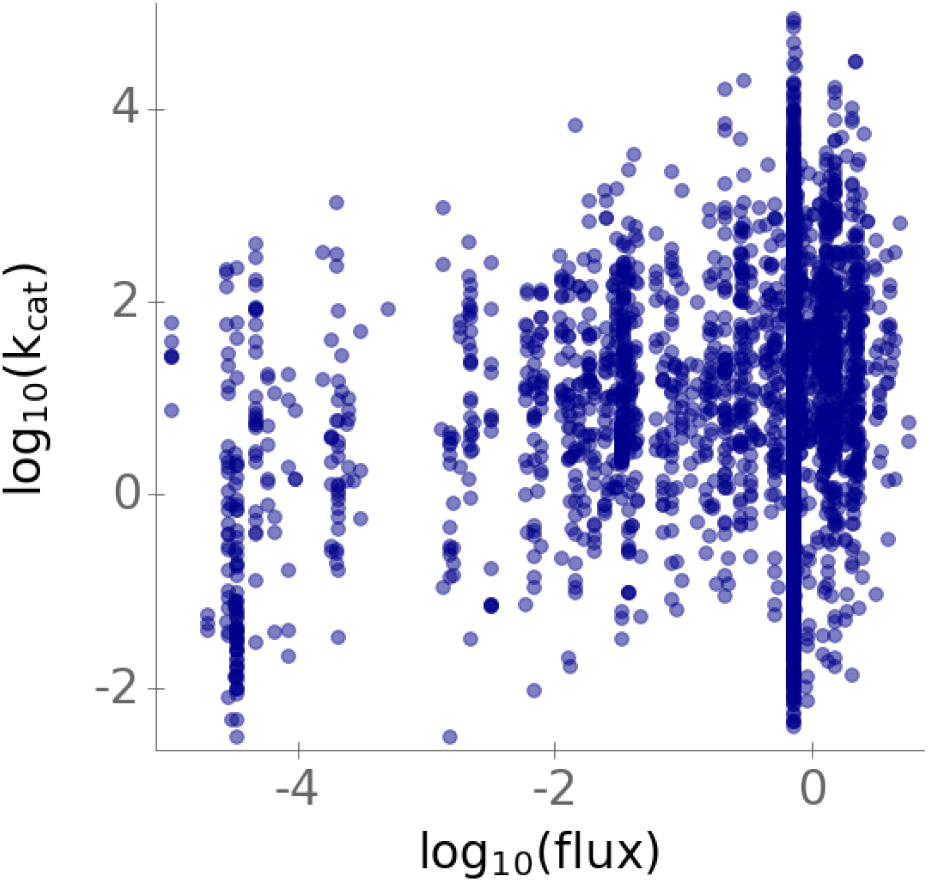
Predicted reaction fluxes compared to *k*_cat_ values. We obtained either predicted reaction fluxes via parsimonious flux balance analysis (pFBA) or flux variability analysis (FVA) and we plotted these values against the corresponding *k*_cat_ values on a log_10_-scale. The plot contains 4271 data points from our training and test set.

